# Global analysis reveals complex demographic responses of mammals to climate change

**DOI:** 10.1101/2019.12.16.878348

**Authors:** Maria Paniw, Tamora James, C. Ruth Archer, Gesa Römer, Sam Levin, Aldo Compagnoni, Judy Che-Castaldo, Joanne M. Bennett, Andrew Mooney, Dylan Z. Childs, Arpat Ozgul, Owen R. Jones, Jean H. Burns, Andrew P. Beckerman, Abir Patwary, Nora Sanchez-Gassen, Tiffany M. Knight, Roberto Salguero-Gómez

## Abstract

Approximately 25 % of mammals are threatened globally with extinction, a risk that is amplified under climate change^1^. Persistence under climate change is determined by the combined effects of climatic factors on multiple demographic rates (survival, development, reproduction), and hence, on population dynamics^2^. Thus, to quantify which species and places on Earth are most vulnerable to climate-driven extinction, a global understanding of how demographic rates respond to climate is needed^3^. We synthesise information on such responses in terrestrial mammals, where extensive demographic data are available^4^. Given the importance of assessing the full spectrum of responses, we focus on studies that quantitatively link climate to multiple demographic rates. We identify 106 such studies, corresponding to 86 mammal species. We reveal a strong mismatch between the locations of demographic studies and the regions and taxa currently recognised as most vulnerable to climate change^5,6^. Moreover, we show that the effects of climate change on mammals will operate via complex demographic mechanisms: a vast majority of mammal populations display projected increases in some demographic rates but declines in others. Assessments of population viability under climate change therefore need to account for multiple demographic responses. We advocate to prioritise coordinated actions to assess mammal demography holistically for effective conservation worldwide.

---

The *ca*. 6,400 extant mammal species^7^ can be found in virtually all terrestrial and most aquatic habitats^8^. This evolutionary success has been facilitated by the wide range of mammalian life history strategies^9^, which enable them to cope with vastly different climates^10^. These strategies include extreme examples like male semelparity in some Australian marsupials with very short mating seasons^11^ or high behavioral and demographic plasticity in long-lived primates that buffers populations from the negative effects of environmental variation^12^. This tremendous variation in life history strategies can be captured by differences among organisms in their rates and timing of survival, development, and reproduction^13^. It is these demographic rates that determine population growth and thus species persistence^14^. Therefore, understanding the effects of climate drivers on the viability of natural mammal populations requires a simultaneous consideration of multiple demographic rates^2^.

Important efforts have been made in the last decade to increase the amount of comparative data to understand the variation in demographic rates across mammals^4,15^. These data have resulted in the broader availability of open-access demographic data on mammal populations^15,16^ and have produced synthetic demographic knowledge, for instance on lifespan and mortality schedules^4,17^. However, we still lack a holistic understanding of how climate drivers simultaneously affect survival, development, and reproduction in mammals worldwide. Consequently, it is unclear whether research quantifying the response of mammal populations to climatic drivers is available for regions most vulnerable to climate change or for the most vulnerable species. Moreover, the complexity of demographic responses to climate remains unknown for most taxa, even in comparatively well-studied groups such as mammals^3^. These knowledge gaps occur despite an emerging consensus that interactions among demographic rates and biotic and abiotic drivers hinder simplistic projections of persistence under climate change^3,18^. For instance, a negative effect of climate on a specific demographic rate does not necessarily cause a population to go extinct, when another demographic rate responds positively to climate, or when population dynamics are mediated by density-dependent feedbacks^2,19^. Consequently, it is vital for demographic research to synthesize available knowledge in how mammalian populations respond to climate drivers given the accelerated loss of mammal species^7^.

Here, we synthesise our understanding regarding where, which, and how mammal populations respond to climate. We conducted a rigorous review of literature linking multiple demographic rates to climatic drivers, thus caputring the complexity of demographic responses, on 5,856 mammal species with available life-history information^20^. We then linked data from the literature review to information on ecoregion and species’ vulnerability to climate change^1,5,21^ to explore (i) whether mammal demographic studies are conducted in ecoregions that are most vulnerable to projected increases in temperature extremes (Q1: *Where*?)^5^; (ii) whether demographic responses to projected changes in climate reflect species’ extinction risk as determined by the IUCN Red List status of mammals (Q2: *Which species?*); and through which demographic processes projected changes in climate may show negative and/or positive effects on populations (Q3: *How?*).

We extracted information on climate-demography relationships from 106 studies, for a total of 86 species, that quantified simultaneous responses to climate in at least two different stage- or age-specific demographic rates. These studies span 14 biomes, with the exception of tropical and subtropical coniferous forests and mangroves (Fig. S1). Overall, more studies assess only the direct effects of precipitation (n = 46) than the direct effects of temperature (n = 11) (Fig. S2); and in 19 of the 106 studies, only indirect effects are assessed via global indices such as the North Atlantic Oscillation (NAO) or El Niño–Southern Oscillation (ENSO). Few studies (10 %) test how different climatic drivers interact with one another, approximately half (55 %) test for the effects of density dependence on demographic rates, and an additional 20 % test for interactions with non-climatic drivers other than population density (e.g. predation, food availability). These omissions may bias estimates of population viability as population dynamics are typically driven by compound effects of interacting climatic and non-climatic drivers^18^, which are projected to become more extreme under climate change^22^.

Our synthesis reveals that few demographic studies are conducted in ecoregions that are both most biodiverse and most vulnerable to climate change. Overlaying the coordinates of the center of each studied population’s range with geographic information on the globally most biodiverse (G200) ecoregions^23^, we find that 41 out of the 106 demographic studies were conducted in one of the G200 ecoregions (Fig. 1). However, only 13 of these studies assess the demographic effects of temperature increases, which, unlike precipitation, is projected to become more extreme in all G200 ecoregions^5^. In addition, no study has examined the responses of different demographic rates in ecoregions with the highest vulnerability scores (e.g., the Central Congo Basin; darkest red in Fig. 1); and only one study, which includes three primate species^12^, assesses temperature effects in relatively highly vulnerable G200 ecoregions. Primates have been shown to buffer the negative effects of climate change via their high behavioral and physiological flexibility^12^. This flexibility may explain why the primate demographic rates were not affected by temperature. In the remaining studies in G200 ecoregions, temperature has positive as well as negative or shows no effects on demography (Fig. 1 insert). This might indicate that the studies did not capture the temperature extremes that are currently occurring in these regions and are expected to increase in frequency in the future. Thus, in addressing “Q1: *Where*?”, our synthesis highlights an urgent need for research on holistic mammal climate-demography relationships in the ecoregions most vulnerable to climate change. Many of these ecoregions also face strong pressures on biodiversity from direct human activities^24^, which are likely to interact with climate change to threaten populations^22^.

**Figure 1:**
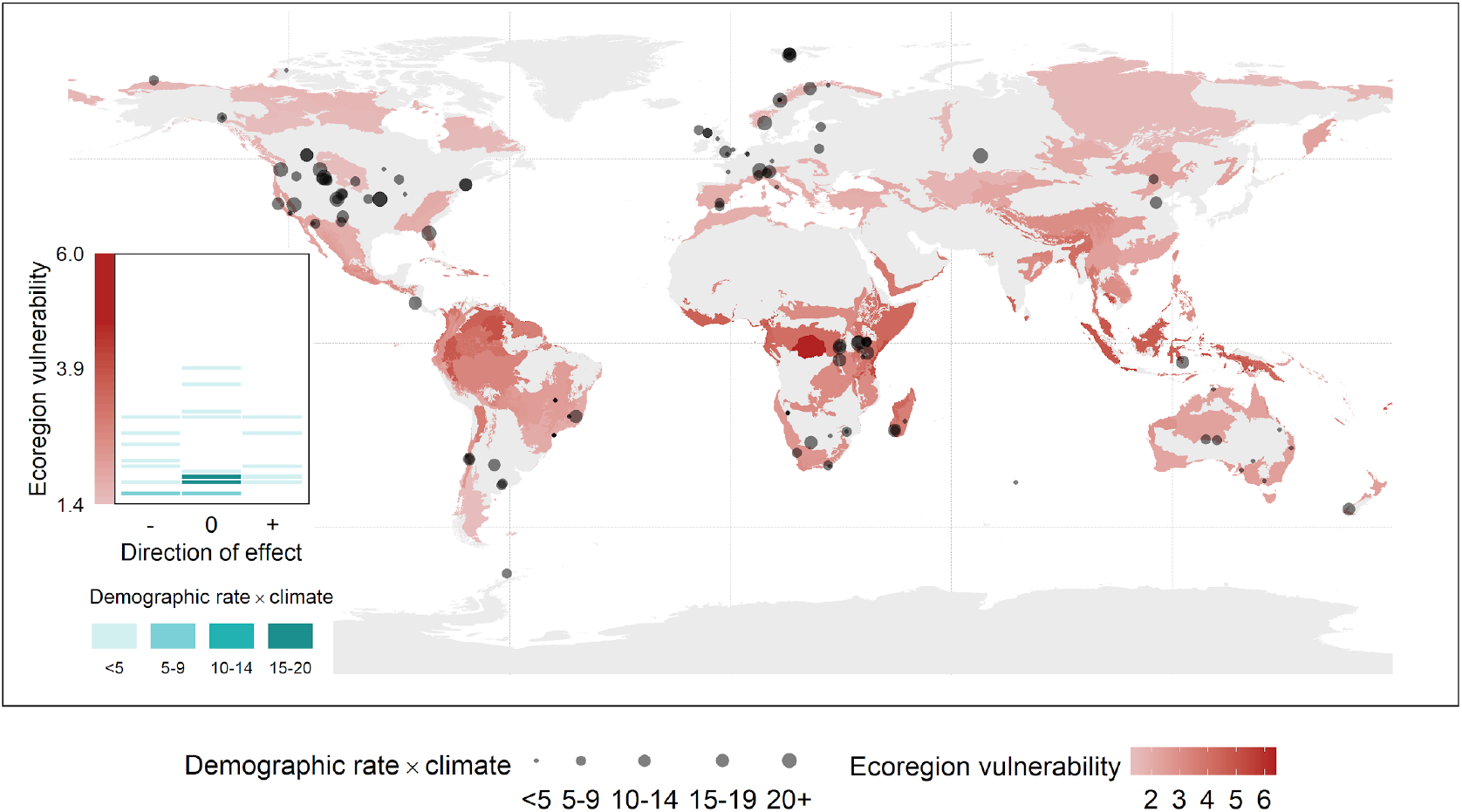
Global distribution of 106 mammal studies (grey points) that have comprehensively assessed demographic responses to climatic drivers across the species’ life cycles. Point size indicates the number of relationships between climatic drivers and stage- or age-specific demographic rates (survival, development, and/or reproduction) assessed. The red-scale background on the map indicates projected climate-change vulnerability for the most biodiverse (G200) ecoregions, with redder colors indicating a higher increase in extreme-temperature events compared to historical conditions. The left insert shows the number of demographic rates decreasing (−), not changing (0), or increasing (+) under increasing temperatures as a function of ecoregion vulnerability. Green shading on the insert indicates the total number of demographic rates linked to temperature in each ecoregion vulnerability level.

In addition to an ecoregion bias, demographic analyses have taxonomic bias. We show that studies linking multiple demographic rates to climatic drivers are primarily performed in regions with a relatively low mammal richness^8,25^ and on species that are not currently vulnerable to climate change (Fig. 2), based on IUCN classifications. Indeed, the IUCN has identified at least 17 % of listed vertebrates to be sensitive to climate change, *i.e.*, decreasing in numbers or losing habitat under changes in temperature and precipitation regimes due to elevated atmospheric CO_2_ levels^26^. Our synthesis reveals that only 4 % of all mammals assessed as climate sensitive by the IUCN have detailed studies linking demography to climate (*i.e.*, 13 % of studies we assessed), allowing this threat to be understood and potentially mitigated through conservation. Interestingly, the proportion of demographic rates per study that will decline under projected changes in climatic drivers (0.31, ± 0.10 S.E.), as assessed in the respective papers or in our analyses, is highest for species that have been flagged by the IUCN as climate sensitive. However, this proportion is followed closely by species for which climate change is not considered a threat by the IUCN (Fig. 2 insert). Therefore, in answering “Q2: *Which species?*”, we highlight the need for future research to prioritise demographic studies for climate-sensitive and threatened mammal species. On the other hand, given that a large number of mammals not considered climate-sensitive by the IUCN may actually show strong negative demographic responses to climate change (Fig. 2), these results also support the need for current IUCN efforts to re-evaluate the importance of climate as an extinction threat to mammals^6^.

**Figure 2.**
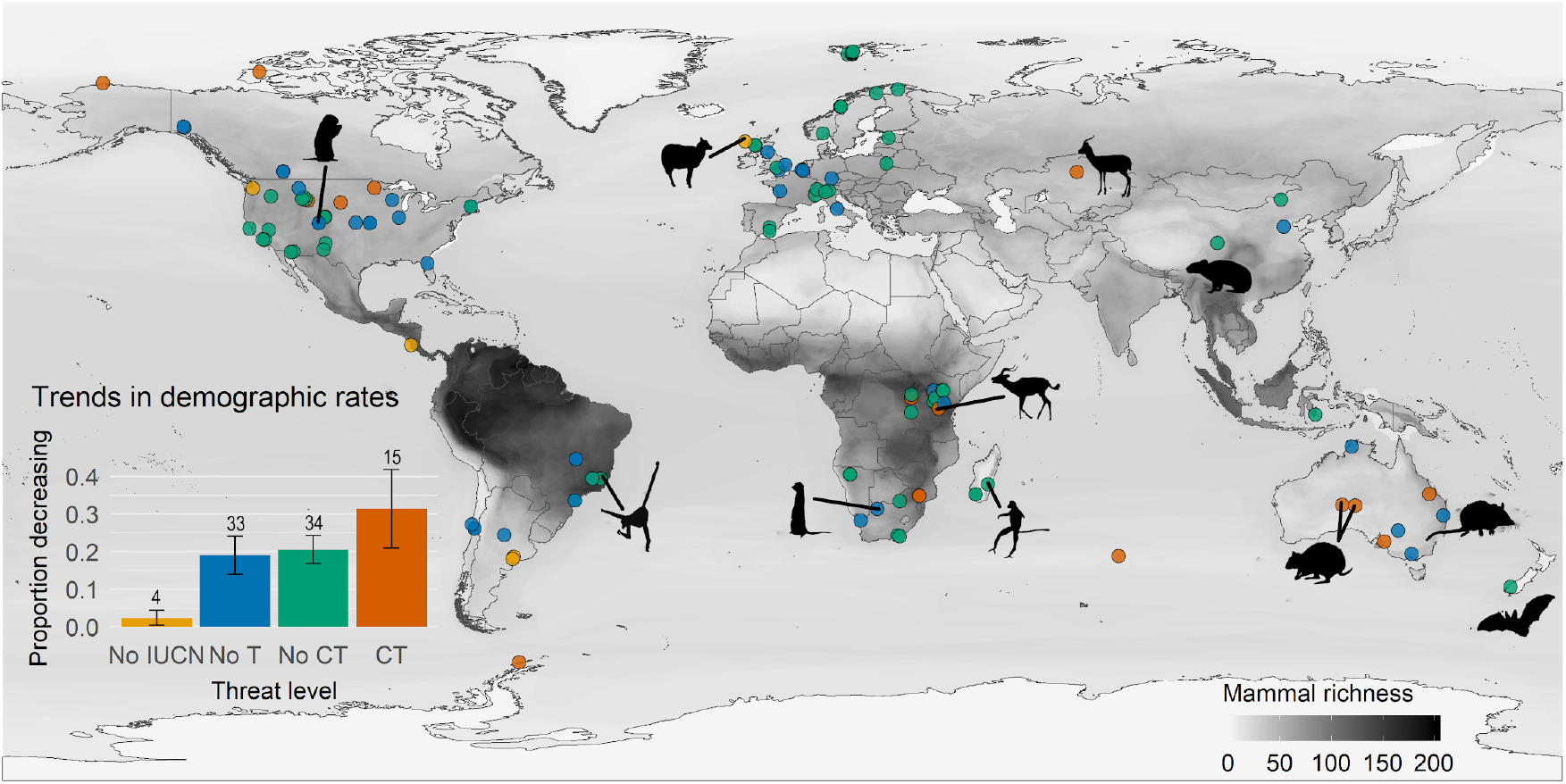
Global distribution of mammals (points) with available information on climate-demography relationships. Point and bar colors indicate levels of threat assessment by the IUCN (No IUCN - species not assessed; No T - species assessed and currently faces no threats; No CT - species assessed and faces threats but climate change is not considered a threat; CT - climate change is considered a threat). Darker background on the map indicates higher mammal richness (number of species). Bottom-left insert displays the mean proportion of demographic rates per studied mammal population ± S.E. (error bars) that will decrease at different magnitudes under projected climate change in different IUCN threat assessment categories. Total number of populations with at least one decreasing rate per threat level are indicated above the bars. Species highlighted in Figure 3 are mapped here using silhouettes.

Across the reviewed studies, multi-directional demographic responses to climate are prevalent. Only eight of the 106 studies report unidirectional (all positive) responses of demographic rates to climatic drivers, while 11 studies find no effect of climate on any demographic rate (Fig. S3). For the vast majority of species, the direction of observed (79 %) and projected (75 %) demographic responses to climate vary depending on the demographic rate or stage/age being considered and on interactions among climatic and non-climatic drivers, with interactions often mediated by density feedbacks (Fig. 3; Fig. S3). For instance, impalas (*Aepyceros melampus*), which the IUCN characterises as threatened by drought (Table S1), may show positive or negative responses in survival and reproductive success under rainfall scarcity (Fig. 3) depending on the seasonal patterning of rainfall and population density^27^. Similarly, meerkats (*Suricata suricatta*), which currently face no threats according to the IUCN, show nonlinear, *i.e.*, both positive and negative, responses to precipitation across demographic rates due to social interactions and density feedbacks^28^. Therefore, as a cooperative breeder, meerkats may be vulnerable to increases in seasonal climatic extremes that decrease group densities^2^. Such complex demographic responses make it challenging to project species’ fates under climate change because the future of populations cannot be accurately determined from single demographic rates^3,19^. Optimistically, our results suggest that complexity of demographic responses may buffer populations from adverse climate effects^29^ (Fig. 3 insert). Therefore, despite the challenges involved in collecting long-term demographic parameters across the entire life cycle^6^, the mechanistic insights gained from such parameters will be invaluable to understand the drivers of biodiversity loss under climate change^3^.

**Figure 3.**
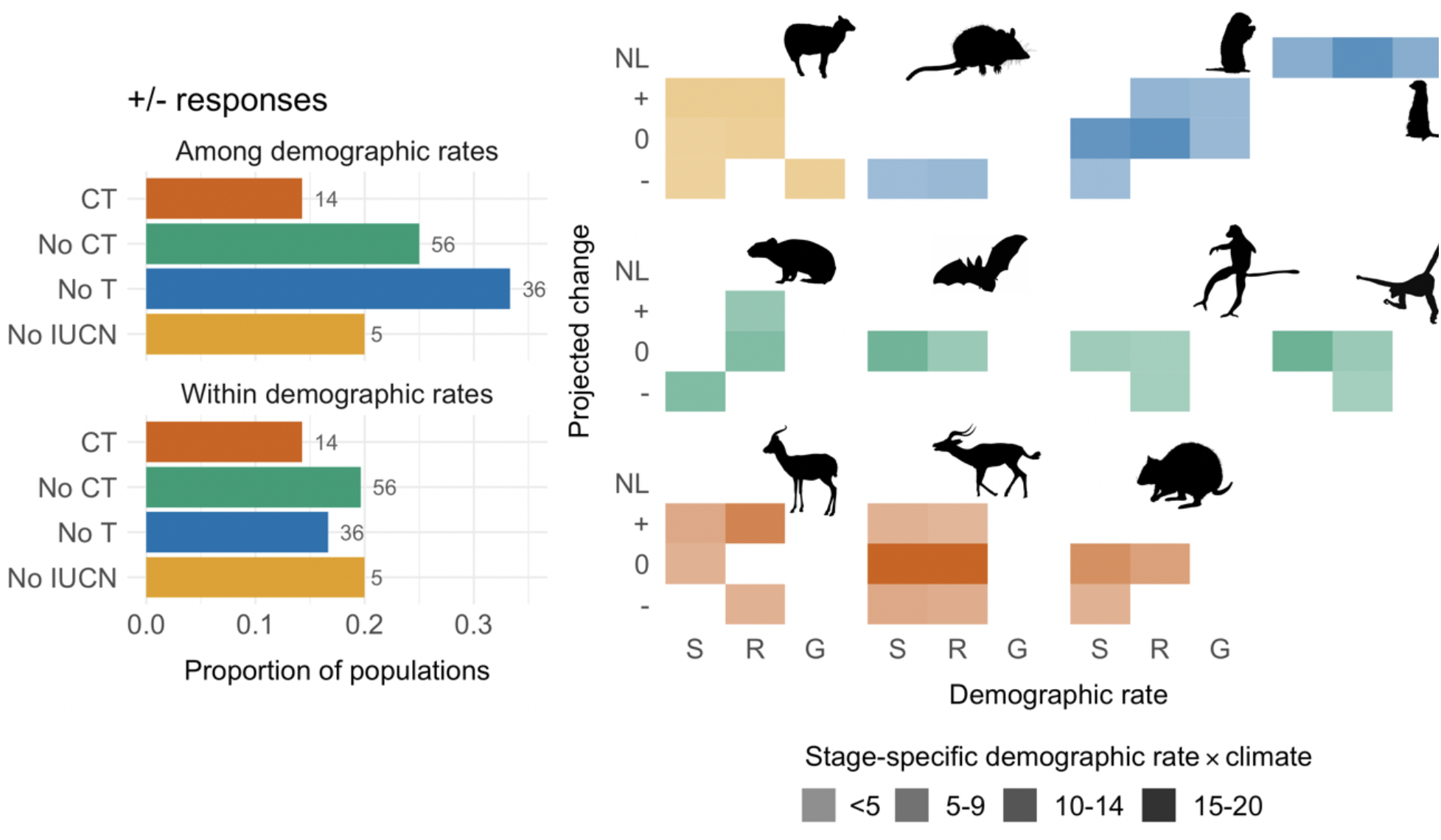
Summary of responses of demographic rates under projected changes in climate across IUCN threat categories (left panel). The proportion of studied populations (out of total number indicated) is shown where the same (within) demographic rate is projected to increase or decrease (+/−) depending on the age/stage modeled; or where a positive response in one rate but negative in another rate (among) are projected. Categories include No IUCN - species not assessed; No T - species assessed and currently faces no threats; No CT - species assessed and faces threats but climate change is not considered a threat; CT - climate change is considered a threat). Detailed responses for 11 example species highlighting the full spectrum of responses are shown in the right panel. Demographic rates include survival (S), probability of reproducing and reproductive output (R), and growth and development (G), which can show only positive (+), only negative (−), nonlinear (NL; both positive and negative), or no (0) responses in the future. From top left to bottom right, the species include Soay sheep (*Ovis aries*), agile antechinus (*Antechinus agilis*), yellow-bellied marmot (*Marmota flaviventer*), meerkat (*Suricata suricatta*), pika (*Ochotona curzoniae*), long-tailed wattled bat (*Chalinolobus tuberculatus*), Milne-Edwards’s sifaka (*Propithecus edwardsi*), northern muriqui (*Brachyteles hypoxanthus*), Saiga antelope (*Saiga tatarica*), impala (*Aepyceros melampus*), and black-flanked rock-wallaby (*Petrogale lateralis*).

By focusing on studies that have assessed several demographic responses to climate, we necessarily limited the number of taxa in our review. In fact, we identified at least 111 more studies on 68 additional species that only assessed climatic effects on single demographic rates. We stress here that we do not question the validity of such studies when population dynamics can be accurately predicted from the changes in one key demographic rate. However, population responses to climate are typically determined by the covariation among multiple demographic rates, which itself is often mediated by a myriad of interacting biotic and abiotic factors, e.g.,^18,19^. In our review, 13 studies assess the effects of climate on population growth rates in addition to underlying demographic rates (Fig. S3, Table S1). These examples show that population responses are not readily predictable from a single demographic rate when multiple climatic drivers and their interactions with biotic drivers affect demography, e.g.,^30^. By revealing the complexity of demographic responses to climate, our synthesis emphasises that projecting population size and structure under climate change requires a complete understanding of demographic processes for most taxa. Therefore, in addressing “*Q3: How?*”, we urge for more studies on climate effects across the whole life cycle of populations.

Mammals are key ecosystem engineers, frequent apex predators, and providers of important ecosystem services^e.g., 31,32^. Future dynamics of mammal populations can therefore determine overall ecosystem change^33^. Our current mechanistic knowledge on mammal responses to climate change would benefit from strategic studies that fill important knowledge gaps. Along with recent calls for a renewed global effort to collect natural-history information^3^, we advocate for a coordinated effort to collect and model demographic responses to climate across the entire life cycle of species, particularly in vulnerable ecoregions such as moist forests in the Congo Basin or mangroves in Madagascar.

## METHODS

### Literature review

We obtained scientific names of all 5,856 mammal species with available life-history information from the Amniote database^20^. For each species *i*, we searched SCOPUS for studies (published before 2018) where the title, abstract, or keywords contained the following search terms:

*Scientific species name*_*i*_ AND (demograph* OR population OR life-history OR “life history” OR model) AND (climat* OR precipitation OR rain* OR temperature OR weather) AND (surv* OR reprod* OR recruit* OR brood OR breed* OR mass OR weight OR size OR grow* OR offspring OR litter OR lambda OR birth OR mortality OR body OR hatch* OR fledg* OR productiv* OR age OR inherit* OR sex OR nest* OR fecund* OR progression OR pregnan* OR newborn OR longevity).

We used the R package *taxize*^*34*^ to resolve discrepancies in scientific names or taxonomic identifiers and, where applicable, searched SCOPUS using all scientific names associated with a species in the Integrated Taxonomic Information System (ITIS; http://www.itis.gov). From any study containing these general search terms, we extracted information on demographic-rate-climate relationships only if the study linked at least two different demographic rates (*i.e.*, survival, development/growth, or reproduction) to a climatic driver (*i.e.*, any direct or indirect measure of temperature or precipitation). In order to focus on robust climate-demography relationships, the response of a demographic rate to a climatic driver had to be quantified using statistical methods, *i.e.*, qualitative or descriptive studies were not included. In addition, for this review, we only considered studies on natural populations of terrestrial mammals, or partially terrestrial mammals (e.g., polar bears), because initial results showed that there were only few climate-related population studies on aquatic mammals, which considered distinct climatic drivers (e.g., sea surface temperatures or ocean circulation indices), lacked future projections, and were not easily assigned to specific ecoregions.

From all studies quantitatively assessing climate-demography relationships, we extracted the following information:

a. Geographic location - The center of the study area was always used. If coordinates were not provided in a study, we assigned coordinates based on the study descriptions of field sites and data collection.
b. Terrestrial biome - The study population was assigned to one of 14 terrestrial biomes^21^ corresponding to the center of the study area. As this review is focused on general climatic patterns affecting demographic rates, specific microhabitat conditions described for any study population were not considered.
c. Climatic driver - Drivers linked to demographic rates were grouped as either local precipitation & temperature indices or global indices (e.g., ENSO, NAO). The temporal extent (e.g., monthly, seasonal, annual, etc.) and aggregation type (e.g., minimum, maximum, mean, etc.) of drivers was also noted.
d. Demographic rate modeled - To facilitate comparisons, we grouped the demographic rates into either survival, reproductive success (*i.e.*, whether or not reproduction occurred), reproductive output (*i.e.*, number or rate of offspring production), growth (including stage transitions), or condition that determines development (i.e., mass or size).
e. Stage or sex modeled - We retrieved information on responses of demographic rates to climate for each age class, stage, or sex modeled in a given study.
f. Driver effect - We grouped effects of drivers as positive (*i.e.,* increased demographic rates), negative (*i.e.*, reduced demographic rate), no effect, or nonlinear (e.g., positive effects at intermediate values and negative at extremes).
g. Driver interactions - We noted any density dependence modeled and any non-climatic covariates included in the demographic-rate models assessing climatic effects.
h. Future projections of climatic driver - In studies that indicated projections of drivers under climate change, we noted whether drivers were projected to increase, decrease, or show nonlinear trends. For studies that provided no information on climatic projections, we quantified projections as described in *Climate-change projections* below (see also `climate_change_analyses_mammal_review.R`).

A full list of extracted studies and a more detailed description of the extraction protocol can be found in the Supporting Information (Table S1). We note that the multitude of methodological approaches used to study demographic responses (e.g. correlation analyses, structured demographic models, individual-based models) renders a meta-analytical approach impractical.

### Ecoregion vulnerability to climate change

We assessed the vulnerability of global ecoregions to climate change following Beaumont and colleagues^5^, who provided a quantitative measure of the sensitivity of ecoregions to climate change. The aforementioned study assessed the likelihood that, by 2070, the “Global 200”, *i.e.*, 238 ecoregions of exceptional biodiversity^23^, would regularly experience monthly climatic conditions that were extreme in 1961–1990. To characterise ecoregions vulnerable to increases in temperature extremes, we first matched the geographic locations of the studied mammal populations to the geographic extent of the G200 ecoregions using the *Intersection* function in QGIS^35^. We then characterised temperature vulnerability of the G200 ecoregions that contained the studied mammal populations using the weighted average minimum monthly distance in temperatures (under the A2 climate model ensemble) from the mean of the 1961-1990 baseline^5^. The higher the distance, the more vulnerable an ecoregion. Lastly, to assess a potential mismatch in demographic studies and ecoregion climate vulnerability (Q1: *Where*?), we quantified the proportion of positive, negative, nonlinear, or no-effect responses of demographic rates to any local temperature variable in each G200 ecoregion. We did not perform this assessment for precipitation, as precipitation extremes were not projected to increase at an ecoregion level ^5^.

### IUCN status of species

To assess whether demographic responses to projected changes in climate (see below) are in agreement with the International Union for Conservation of Nature and Natural Resources (IUCN) Red List status of mammals (Q2: *Which species?*), we obtained IUCN assessments (including threats) for all species identified in the literature review. We used the R package *rredlist* to access the IUCN Red List database and extract available information on whether the species are listed in the database, and, if so, what status they are assigned to and whether climate change is listed as an existing or potential threat.

### Climate-change projections

For studies which did not report on “*future projections of climatic driver*” (70% of studies), we quantified such future projection for climatic variables that depicted direct precipitation and temperature measures. For global indices such as ENSO or NAO, future projections could not be obtained (with the exception of the ones explicitly discussed in a given study), as such projections are either lacking or extremely complex and uncertain^36–38^. All analyses can be replicated using the R script `climate_change_analyses_mammal_review.R`. To project future changes in temperature and precipitation, we obtained monthly average temperatures and rainfall data as well as maximum and minimum monthly temperatures from 1979-2013 for all relevant study locations using *climatologies at high resolution for the earth’s land surface areas* (CHELSA)^39^. We averaged these historical climate records for each month and calculated standard deviation across months, which we could then link to studies that assessed the effects of such deviations. We also obtained monthly projected values of theses variables averaged from 2041 to 2060. We obtained values from five diverging climate models that used different methods for projections assuming a representative concentration pathway of 4.5 W/m^2^ (http://chelsa-climate.org/future/). For each relevant study that assessed averages or deviations in precipitation or temperature (or minimum/maximum temperatures), we quantified whether a given driver was projected to either increase or decrease (95 % CI across the five projection models did not cross historical values) or show no change (95 % CI crossed historical values). From this information, we then determined whether a demographic rate would decrease (e.g., where a rate has a positive response to precipitation and precipitation projected to decrease) or increase (e.g., where a rate has a positive response to precipitation and precipitation projected to increase). Unless explicitly stated otherwise in a study, we assumed that demographic rates that were not affected by a climatic variable would not change in the future, and ones that showed nonlinear responses would also likely show nonlinear responses in the future^2,40^.

## Acknowledgements

This work was supported by the working group proposal “sAPROPOS: Analysis of PROjections of POpulationS” funded to RS-G by the German Centre for Integrative Biodiversity Research (iDiv). MP was supported by an ERC Starting Grant (33785) and a Swiss National Science Foundation Grant (31003A_182286) to AO; and by a Spanish Ministry of Economy and Competitiveness Juan de la Cierva-Formación grant FJCI-2017-32893. RS-G was also supported by a NERC grant (R/142195-11-1). We thank the Alexander von Humboldt foundation (award to TMK) that supported a retreat to write a first version of this manuscript.

## Author contributions

MP, TJ, GR, and RS-G devised the overall manuscript. MP and TJ designed the literature review protocol, which was then implemented by MP, TJ, GR, CRA, SL, AM, JC, NSG, JMB, and AP. The climatic data were derived by AC. The first draft of the manuscript was written by MP and RS-G, and all co-authors contributed to the final manuscript. See Table S2 for further specifics regarding task contributions.

## Competing interests

The authors declare no competing interests.

## Data availability

The data that support the findings in this study are available in the Supplementary Online Materials, Table S1.

## Code availability

The code that supports the findings in this study is available in the Supplementary Online Materials, `climate_change_analyses_mammal_review.R`.

**Supplementary Information** is available in the online version of the paper.

## Online Supplementary Material

**Figure S1.**
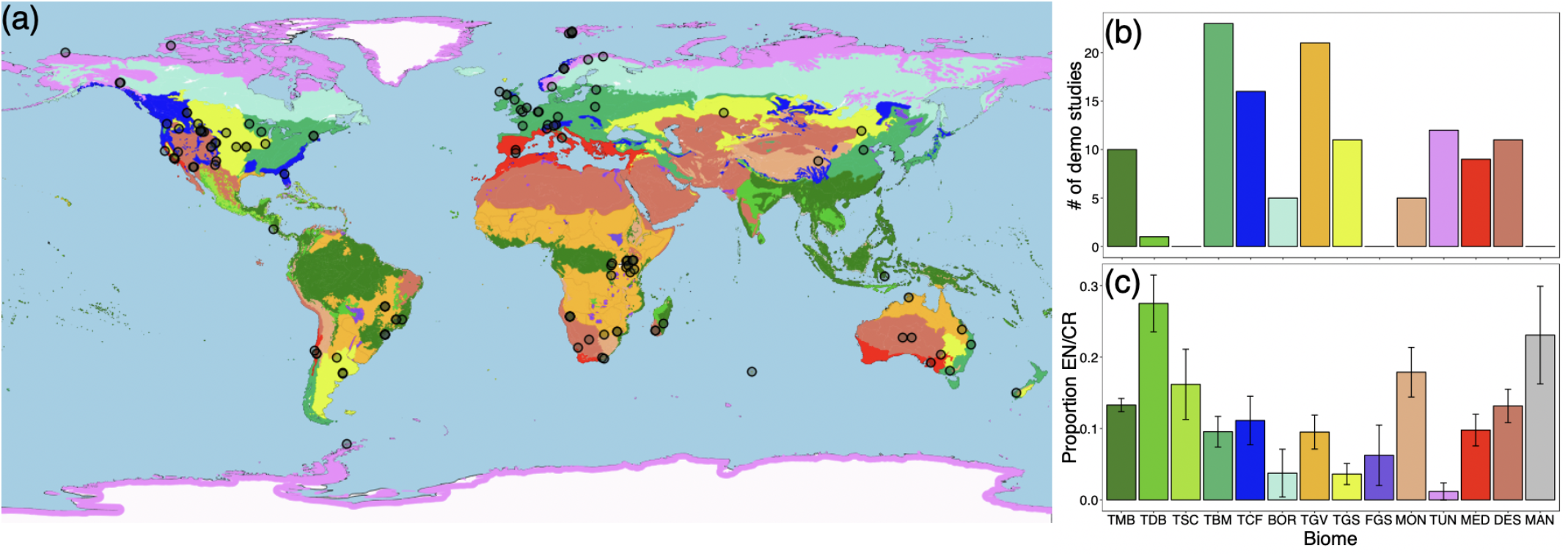
(a) Geographic location of the 106 publications examined in this study that have explicitly evaluated the effect of climate change on mammal population dynamics. (b) Representation of these studies and (c) proportion of mammal species that are endangered (EN) or critically endangered (CR; IUCN Red List of Threatened Species) aggregated by terrestrial biome. TMB: Tropical and Subtropical Moist Forests; TDB: Tropical and Subtropical Dry Forests; TSC: Tropical and Subtropical Coniferous Forests; TBM: Temperate Broadleaf and Mixed Forests; TCF: Temperate Coniferous Forests; BOR: Boreal Forests/Taiga; TGV: Tropical and Subtropical Grasslands, Savannas, and Shrublands; TGS: Temperate Grasslands, Savannas, and Shrublands; FGS: Flooded Grasslands and Savannas; MON: Montane Grasslands and Savannas; TUN: Tundra; MED: Mediterranean Forests, Woodlands, and Shrubs; DES: Deserts and Xeric Shrublands; MAN: Mangrove. Plot in (c) depicts the average (± SE) proportion across polygons classified as a given biome and standardised by polygon area.

**Figure S2.**
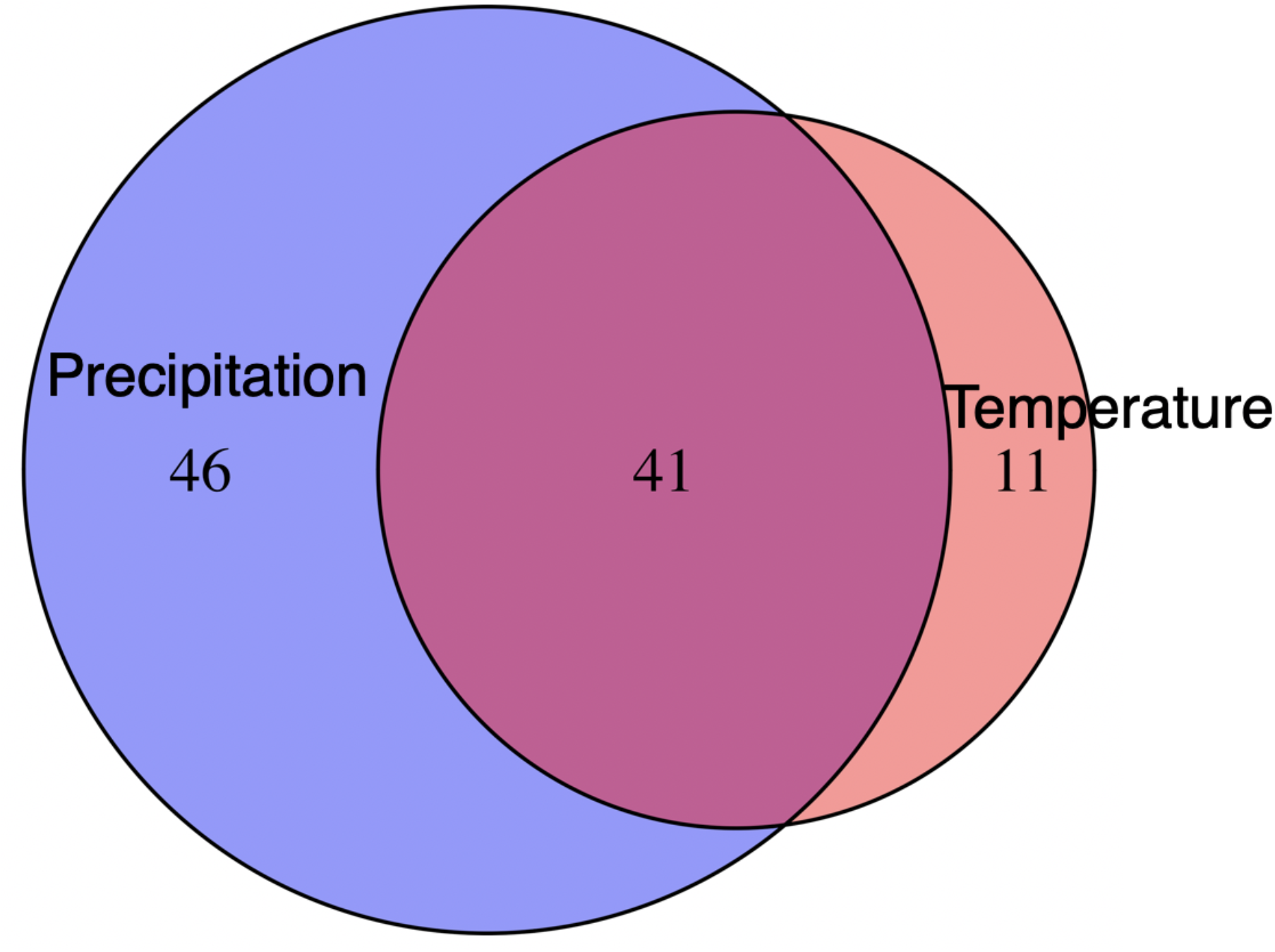
Venn diagram representing (area) the number of studies included in our literature review that explicitly linked mammal demographic responses to precipitation (cyan), temperature (red) or both (purple).

**Figure S3.**
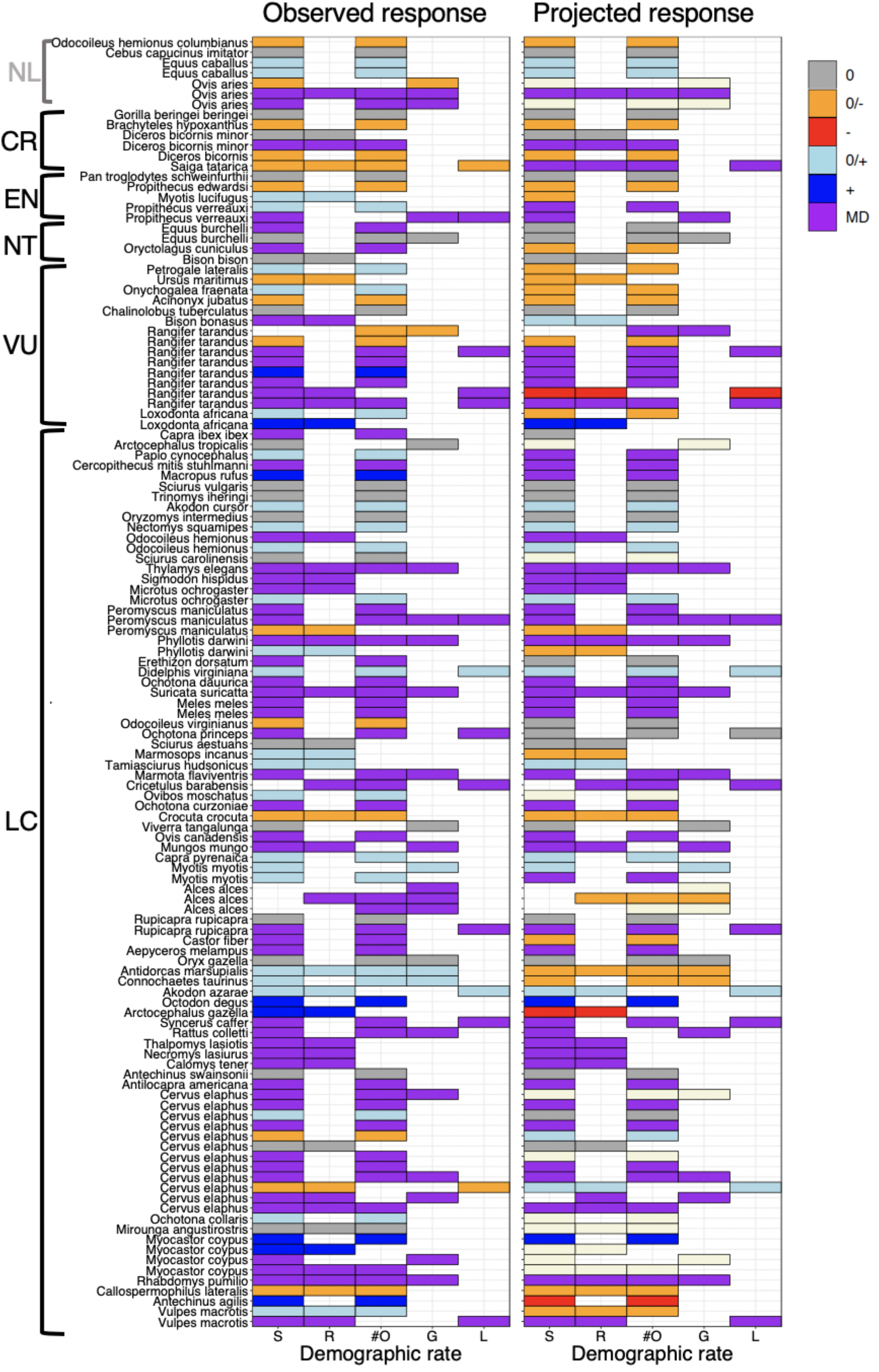
Observed (extracted from demographic studies) and projected (see *Climate-change projections* in Methods) responses of demographic rates for all mammal species reviewed. Species are sorted by the IUCN threat categories: least concerned (LC), vulnerable (VU), near-threatened (NT), endangered (EN), critically endangered (CR). The topmost species have not been assessed (NL) by the IUCN. Demographic rates include survival (S), probability of reproducing (R), reproductive output (#O), growth and development (G), and population growth (L), which increase (+), decrease (−), or show multidirectional (MD; increase for one life-cycle stage or range of climate and decrease for another) or no (0) responses. Demographic rates for which future changes under projected climate change could not be obtained because these rates were modelled as functions of global indices (e.g., ENSO) that are difficult to project are plotted in beige (right plot). Repetition of species names occurs because several publications assessed climate-demography relationships for some species (e.g. *Ovis aries*).

**Table S1.** List of all extracted information on demographic studies that assessed responses to climatic drivers in at least two vital rates. Available as a text file at XXX

## Detailed Extraction Protocol and Data Description

### Protocol Summary

Data were extracted from papers by a team of digitisers (see Table S2), each of whom worked independently on a randomly assigned collection of species. A formatted data-sheet was provided to facilitate consistent and standardised data extraction. Once individuals had collected data, the resulting dataset was error checked in a number of ways. For example, digitisers randomly checked 10 % of papers in the database entered by colleagues, to ensure that outputs from two different digitisers were consistent. Error-checkers also ensured that there were no duplicated manuscripts recorded (this could conceivably happen if a paper modelled more than one species and digitisers extracted data for all species studied in a particular manuscript) and also that all data were entered in a standardised format. Here, we describe all of the data that were collected, and how each item of data was defined.

### Data Description

#### 1. Location data

##### a. Latitude and longitude

The latitude and longitude of a particular study site (as reported in the manuscript) were recorded in decimal degrees using the WGS84 global projection. Notes were also made on how the location was described in the paper, *i.e.* if the location provided represented the middle of a study site, or how latitude and longitude were calculated for migratory species. If latitude and longitude were not reported in the original manuscript, the digitisers used the verbal description of the study site (e.g. nearest town, center of national park etc. where the study was conducted) to estimate these values. Such an approximation of study location did not affect our analyses and conclusions, which were based on broad-scale ecoregion comparisons and on climate data that were interpolated over a relatively large grid of approximately 1 km^2^.

##### b. Biomes and ecoregions

We obtained georeferenced maps of terrestrial biomes and ecoregions from the World Wildlife Fund^25^. Each location identified in our review could therefore be placed into a biome that consisted of one or more ecoregions, some of which correspond to highly diverse G200 ecoregions. Terrestrial biome categories included: **TMB** – tropical and subtropical moist broadleaf forests; **TDB** – tropical and subtropical dry broadleaf forests; **TSC** – tropical and subtropical coniferous forests; **TBM** – temperate broadleaf and mixed forests; **TCF** – temperate coniferous forests; **BOR** – boreal forests / taiga; **TGV** – tropical and subtropical grasslands, savannas and shrublands; **TGS** – temperate grasslands, savannas and shrublands; **FGS** - flooded grasslands and savannas; **MON** – montane grasslands and shrublands; **TUN** – tundra; **MED** – Mediterranean forests, woodland and scrubs; **DES** – deserts and xeric shrublands; **MAN** – mangroves. Definitions for each of these biomes as well as all ecoregions can be found at http://wwf.panda.org/about_our_earth/ecoregions/ecoregion_list/.

#### 2. Climatic Data

##### a. Climatic Drivers

Climatic drivers were divided into the following categories: **P** - any measure of precipitation; **T** - any measure of temperature; **PT** - measures such as drought or icing that reflect both temperature and precipitation. Some climatic drivers were variables derived from raw measures of precipitation and temperature. These variables were described as in the reviewed papers and include **NAO** - Northern Atlantic Oscillation, **ENSO** - El Niño–Southern Oscillation; **SAM** - Southern Annular Mode; **SOI** - Southern Oscillation Index, **PDSI** - Precipitation and Surface Air Temperature and **PDO** - Pacific Decadal Oscillation. A detailed description of each of the climatic drivers included in the dataset was also recorded, to facilitate error checking and data-standardisation.

##### b. Temporal Aggregation

How climatic data were aggregated in statistical models was recorded, with options being: **D** - daily; **S** - seasonal; **M** - monthly; **A** - annual.

##### c. Aggregation Methods

The method used to aggregate climatic data was recorded with options including **sum** - the sum of all climatic values; **min** - the minimum observed value; **max** - the maximum observed value; **mean** - the average value; **SD** - standard deviation in climatic values; **range** - difference between minimum and maximum observed values; **length** - number of days, or growing degree days.

#### 3. Response Traits

##### a. Demographic rates

The studies that feature in the dataset quantified demographic rates in different ways. Accordingly, we grouped the rates featuring in each paper as being associated with survival, reproductive success, reproductive output, growth/development, condition, or population growth. Here, we outline how we assigned traits from individual studies to each of these classes.

###### Survival

Both mortality rates and survival rates feature in our database. However, to ensure that these rates were comparable between studies we reported the sign of any effect as being appropriate for a measure of survival, *i.e.* an environmental variable that increased mortality risk, was recorded in our dataset as reducing survival.

###### Reproductive Success and Output

Studies quantifying reproduction may have recorded the probability of reproduction, number of offspring, reproductive success, number of litters, birth rate, fecundity, reproductive rate, pregnancy or transition into reproductive state. For the purpose of our analyses, any binary variable that defined whether a reproductive event occurred or not, was recorded as a measure of ***Reproductive Success***, while any measure of how many, or how frequently offspring were produced was classed as ***Reproductive Output***.

###### Growth/Development

Variables that quantified individual growth rates, development or generation time were included as measures of growth.

###### Condition

In some cases condition was quantified explicitly using a species specific parameter, but in other cases mass or body size was measured.

##### b. Stage, State or Sex Modelled

Digitisers recorded which life-stage (i.e. juvenile, adult), sex and state (e.g. individual size for IPMs) was modelled, using the description provided by the authors in the manuscript. If an unstructured population model was used, this was recorded as “unstructured”.

##### c. Direction of effect

Digitisers recorded if the climatic driver has a negative effect on the demographic rate (**neg**), a positive effect (**pos**), a nonlinear effect (**nonlinear**) or no effect **(noe**).

##### d. Duration of Study

The number of years that data were collected was recorded.

#### 4. Model Details

To understand the nature of the models collected in our data-base, for example, how often existing data quantifies interactions between climatic variables, the details of the model were recorded as described below.

##### a. Density Dependence

Digitisers recorded whether data dependence was modelled (binary variable, **yes** or **no**).

##### b. Indirect Effect of Driver

Digitisers recorded if indirect effects, e.g., path analyses, were tested for in the model (binary variable, **yes** or **no**).

##### c. Non-linear Effect of Driver

If a climatic driver had a non-linear effect on the demographic rate, the nature of that effect was described here, with examples including **quadratic**, **lag** or **other**.

##### d. Interaction with Other Climatic Driver(s)

Were interactions considered between climatic drivers (binary variable, **yes** or **no**)?

##### e. Interaction with Other Non-Climatic Driver(s)

Were interactions considered between climatic drivers and other variables not related to climate? Digitisers recorded **yes** or **no**

##### f. Non-Climatic Drivers

Where relevant, a description of the non-climatic driver(s) modelled was recorded as concisely as possible.

##### g. Future Driver Direction

If described in a paper, we noted how the climatic driver modelled was expected to change under current climatic change models. Options included **increase**, **decrease**, **nonlinear**, or **no change**.

**Table S2.**
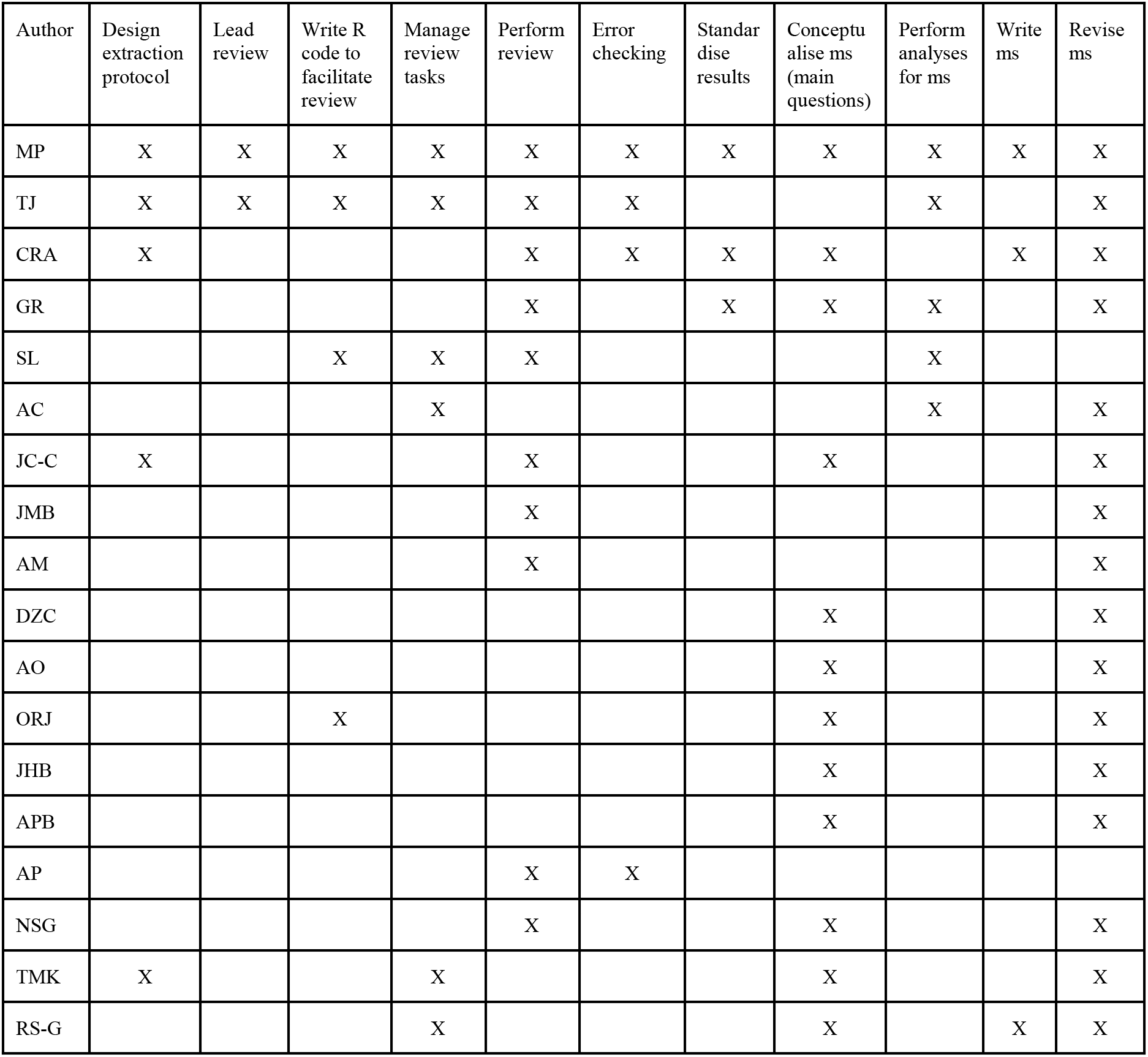
Extended task contribution by each author in this manuscript (ms)

## Additional Sources of Support

This work emanated from the sDiv working work “sAPROPOS” led by RS-G. RS-G was also supported by NERC (NE/M018458/1). JC-C, RS-G, and ORJ. were also supported by an NSF Advances in Biological Informatics grant (DBI-1661342000). TMK was funded by the Helmholtz Recruitment Initiative of the Helmholtz Association and by the Alexander von Humboldt Foundation in the framework of the Alexander von Humboldt Professorship. TJ was funded through “Adapting to the Challenges of a Changing Environment”, a NERC funded doctoral training partnership (ACCE DTP; NE/L002450/1). ORJ was supported by the Danish Council for Independent Research (grant # 6108-00467).

